# Chill coma onset and recovery fail to reveal true variation in thermal performance among populations of *Drosophila melanogaster*

**DOI:** 10.1101/2020.12.10.419341

**Authors:** Hannah E. Davis, Alexandra Cheslock, Heath A. MacMillan

## Abstract

1. Species from colder climates tend to be more chill tolerant regardless of the chill tolerance trait measured, but for *Drosophila melanogaster*, population-level differences in chill tolerance among populations are not always found when a single trait is measured in the laboratory.
2. We measured chill coma onset temperature, chill coma recovery time, and survival after chronic cold exposure in replicate lines derived from multiple paired African and European *D. melanogaster* populations. The populations in our study were previously found to differ in chronic cold survival ability, which is believed to have evolved independently in each population pair.
3. To our surprise, the populations did not differ in chill coma onset temperature and chill coma recovery time in a manner that reflected their geographic origins, even though these traits are known to vary with origin latitude among *Drosophila* species and among the most common metrics of thermal tolerance in insects. The populations did, however, still differ in their ability to survive chronic cold exposure.
4. While it is common practice to measure only one chill tolerance trait when comparing chill tolerance among insect populations, our results emphasise the importance of measuring more than one thermal tolerance trait to minimize the risk of missing real adaptive variation in insect thermal tolerance.

## Introduction

Thermal limits partly determine where insects can live: several measures of chill tolerance correlate strongly with latitudinal (Addo-Bediako, Chown, & Gaston, 2000; J. L. Andersen et al., 2015; Kimura, 2004) and elevational distribution (Gaston & Chown, 1999). While a minority of insects can endure freezing temperatures, most die at temperatures above freezing for reasons unrelated to ice formation. An insect’s ability to survive and function at (relatively) mild cold temperatures is its chill tolerance, and this term can refer to a complex suite of traits: the temperature at which an individual enters a chill coma (a state of temporary paralysis induced by cold), its speed of recovery from chill coma when returned to benign conditions, its ability to withstand cold without injury or death, and its fitness (quantified in any number of ways) after cold exposure (MacMillan, 2019; Overgaard & MacMillan, 2017).

It is becoming increasingly clear that different measures of cold tolerance allow us to see the effects of cold on different organ systems, albeit indirectly. Chill coma onset, for example, involves neuromuscular signal transmission failure. Waves of spreading depolarisation first shut down the central nervous system (Armstrong, Rodríguez, & Meldrum Robertson, 2012; Rodgers, Armstrong, & Robertson, 2010) at the critical thermal minimum (CT_min_; the temperature at which movement becomes uncoordinated), and this is typically followed by muscle membrane depolarisation, and complete paralysis at the chill coma onset temperature (CCO) (M. K. Andersen & Overgaard, 2019).

Insects need to restore ion balance upon rewarming because this balance is lost in the cold; insects susceptible to chilling progressively lose haemolymph ion and water homeostasis while they remain chilled (Koštál, Vambera, & Bastl, 2004; Overgaard & MacMillan, 2017). At permissive temperatures, ion and water homeostasis are typically maintained through active ion pumping in the renal organs – the Malpighian tubules and the rectum – but active transport slows in the cold (Zachariassen, Kristiansen, & Pedersen, 2004). In the cold, active transport cannot compensate for passive ion leak, and as haemolymph Na^+^ and water leak down their own concentration gradients into the gut, K^+^ is concentrated in the remaining haemolymph (Koštál et al., 2004; MacMillan & Sinclair, 2011). High haemolymph [K^+^] causes muscle cell depolarisation, so it is thought that ability to recover K^+^ homeostasis and the degree to which homeostasis is lost in the cold determine an insect’s chill coma recovery time (M. K. Andersen & Overgaard, 2019; Koštál et al., 2004; MacMillan, Findsen, Pedersen, & Overgaard, 2014; MacMillan, Williams, Staples, & Sinclair, 2012). Survival is also linked to the degree to which homeostasis is lost in the cold: high haemolymph [K^+^] is toxic and can lead to chilling injury and death by triggering cellular Ca^2+^ overload (Bayley, Sørensen, Moos, Koštál, & Overgaard, 2020; Bayley et al., 2018; Carrington, Andersen, Brzezinski, & MacMillan, in press; Koštál, Yanagimoto, & Bastl, 2006; MacMillan, Baatrup, & Overgaard, 2015).

Since multiple organ systems are involved, chill tolerance traits can vary independently (Garcia, Littler, Sriram, & Teets, 2020; Gerken, Mackay, & Morgan, 2016) – but in practice, they often don’t. After cold acclimation or rapid cold hardening (i.e. pre-exposure to a sublethal cold stress), multiple chill tolerance traits are often observed to improve (albeit not always to the same extent) (Colinet & Hoffmann, 2012; MacMillan, Andersen, Loeschcke, & Overgaard, 2015; Ransberry, MacMillan, & Sinclair, 2011). Correlations among related traits are also common in nature, and species and populations from colder regions tend to be more chill tolerant overall (i.e. we often see correlations among chill tolerance traits as well as between individual traits and geographic distribution) than species or populations from warmer regions (Addo-Bediako et al., 2000; Sunday, Bates, & Dulvy, 2011). Among *Drosophila* species, CCO correlates strongly with two measures of survival (lethal temperature and lethal time at low temperature), weakly with CCRT, and these traits also correlate with geographical distribution (albeit only weakly, in the case of CCRT) (J. L. Andersen et al., 2015). Within a single species, *Drosophila melanogaster*, populations from colder regions are sometimes, but not always, more chill tolerant based on one or more of these traits. For example, higher latitude Australian populations are more chill tolerant based on CCO, CCRT and cold shock survival (A. A. Hoffmann, Shirriffs, & Scott, 2005; Ary A. Hoffmann, Anderson, & Hallas, 2002; Overgaard, Hoffmann, & Kristensen, 2011). Similarly, in a large study of *D. melanogaster* populations from around the world, temperate populations were more chill tolerant based on CCRT (the only measure used) than tropical populations (Ayrinhac et al., 2004); however, in a separate study of African and European *D. melanogaster* populations, critical thermal minimum was not related to latitude (although it was in a related species, *D. subobscura*) (Gibert & Huey, 2001), and in Japan, there is minimal variation in cold shock survival and CCO among northerly/southerly populations within *Drosophila* species (although species with more northerly distributions are more chill tolerant based on those same measures) (Hori & Kimura, 1998; Kimura, 2004).

Patterns of variation of chill tolerance within *Drosophila* species is further complicated by not knowing how previously-tested population sets are related: CCO, at least, has a very strong phylogenetic signal (Kellermann et al., 2012), which can cloud interpretation if not accounted for. One system for which phylogenetic relationships are known is the set of African and European populations of *D. melanogaster* used by Pool et al. (2017) to study the parallel evolution of chill tolerance traits. The system consists of paired, closely-related populations and a single outgroup (Zambia; *D. melanogaster* originally evolved in the warm, tropical forests of Zambia and Zimbabwe (Mansourian et al., 2018; Pool et al., 2012)). In each pair, one population is derived from flies caught in a relatively cold region, whereas the other is derived from flies caught in a relatively warm region (Figure 1).

**Figure 1.**
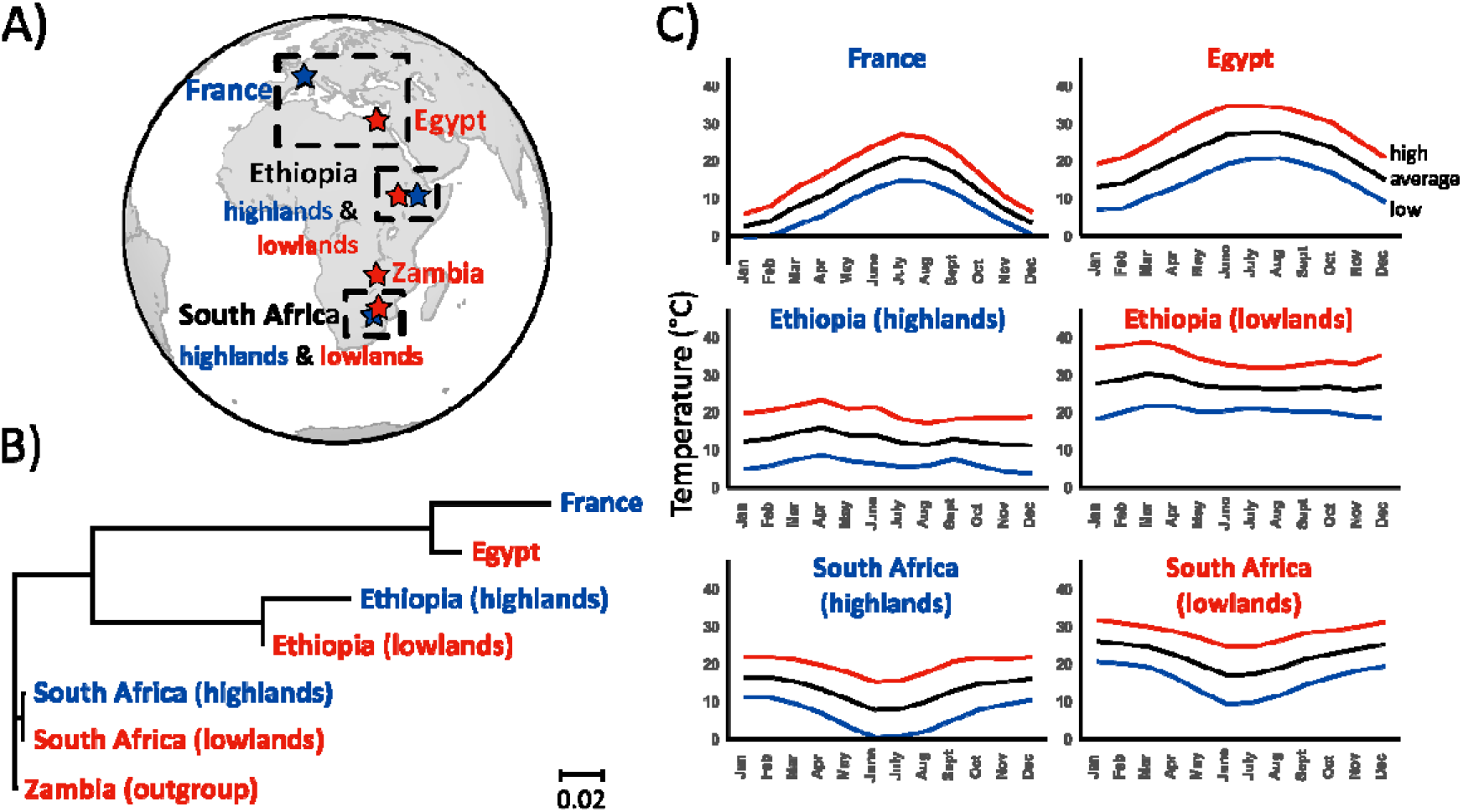
Parallel lines study system. A) Approximate geographic locations where the progenitors of the laboratory populations were collected, with boxes enclosing paired populations. B) Neighbour-joining tree showing the relatedness of the populations, modified from Pool et al. (2017). C) Average monthly highs and lows in the locations where the founder populations of the paired laboratory populations were collected. All climate data was downloaded from climate-data.org.

**Figure 2.**
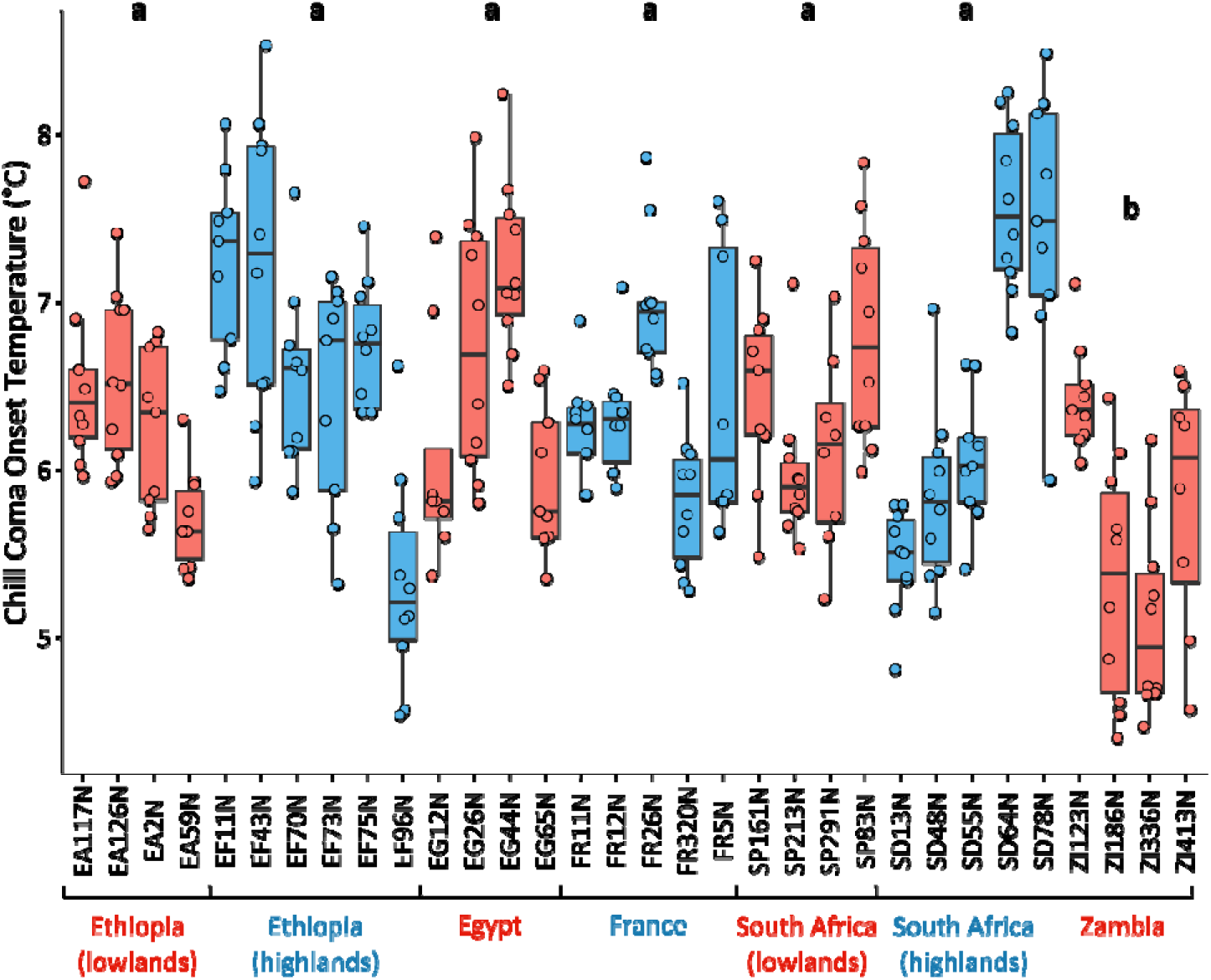
Chill coma onset temperature of all lines. Lines in blue are from relatively cold habitats, whereas lines in red are from relatively warm habitats. The thicker line in the centre of each boxplot represents the median, and the upper and lower hinges represent first and third quartiles, respectively. Whiskers extend to 1.5 times the interquartile range, or to the smallest or largest values if they fall within 1.5 * IQR. Points represent individual flies. Different letters (a or b) indicate a significant difference (p < 0.05) at the population level.

Pool et al. (2017) used a chronic cold exposure assay (4 days at 4°C) to demonstrate that populations from cold regions are more chill tolerant, and concluded that this tolerance likely evolved independently in each cold-region population. This system provides an opportunity to also examine whether and how other chill tolerance traits have evolved in cold vs. warm climates, without the concern that observed chill tolerance in separate regions might be inherited from a common ancestor rather than independently evolved. We thus set out to further characterise these populations using two of the most common chill tolerance measures – chill coma onset temperature (CCO) and chill coma recovery time (CCRT). Because chill tolerance traits often do correlate both with each other and with climate of origin in natural systems, we expected that cold-climate populations would be more chill tolerant than their warm-climate counterparts based on one or both of these traits. To our surprise, the populations differed very little based on either the CCO or CCRT assays – within or among pairs. To ensure that differences in chill tolerance had not simply been lost via lab adaptation, we further characterised survival during a 4 day exposure to 4°C in French and Egyptian lines (the pair previously found to exhibit the greatest difference in recovery after chronic cold exposure (Pool et al., 2017)) and confirmed clear differences in at least one measure of chill tolerance persist in these populations.

## Methods

### Study system

The study system used here (supplied and previously studied by Pool et al. (2017)) consists of seven *Drosophila melanogaster* populations: three pairs of closely-related populations plus one outgroup (Zambia) (Table 1). Within each population pair, one population is derived from flies collected in a relatively warm location (e.g. Ethiopian lowlands), while the other is derived from flies collected in a relatively cold location (e.g. Ethiopian highlands).

**Table 1.**
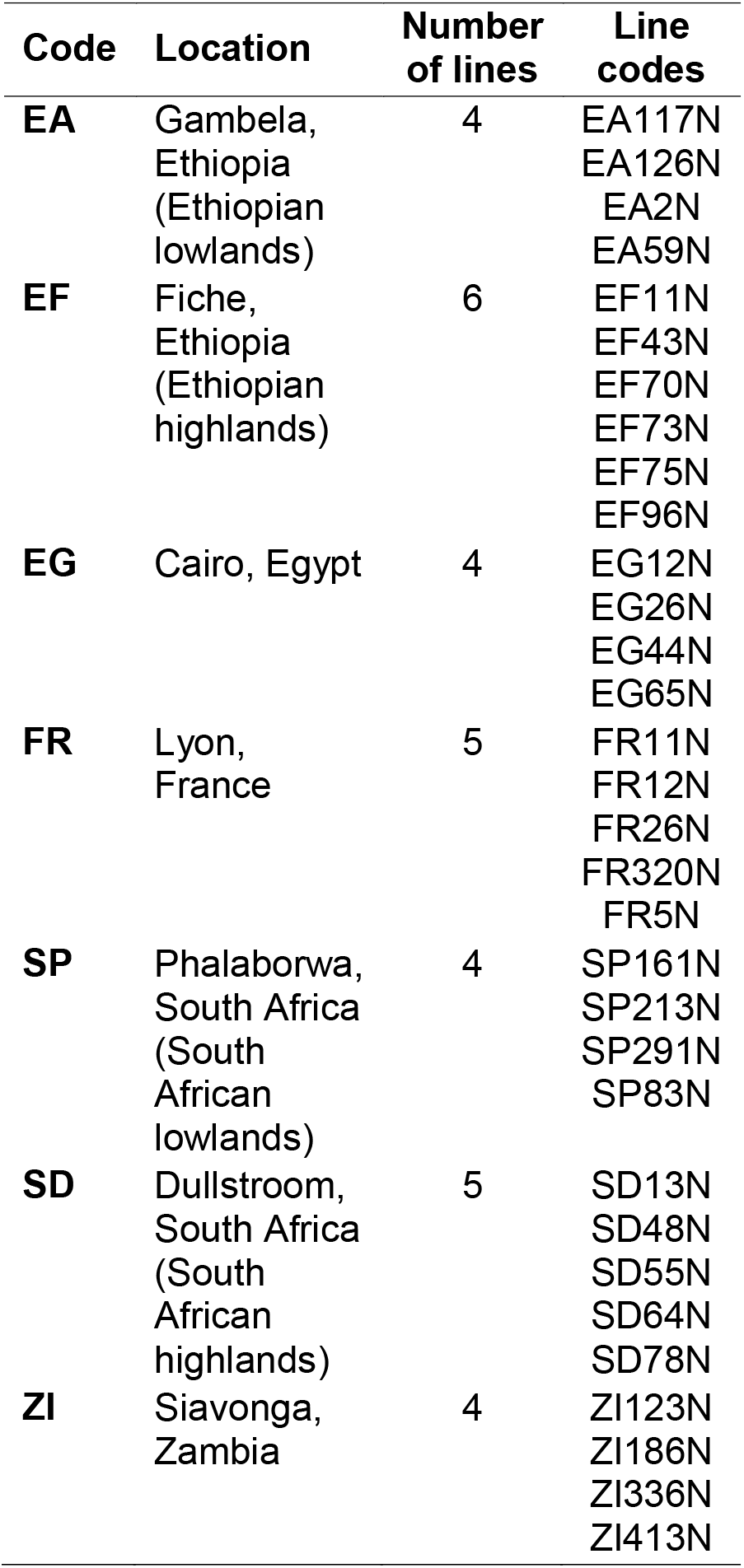
Origins of the *Drosophila melanogaster* populations previously used by Pool et al. (2017). These lab-reared populations comprise multiple isofemale lines, each derived from a female fly captured in the wild in one of seven locations.

To compare populations’ cold tolerance, Pool et al. (2017) exposed flies to 4°C for 96 hours (4 days), returned them to room temperature, and then recorded the number that regained the ability to move or stand within 30 minutes. Within each population pair, the population from the colder climate was significantly more cold tolerant than its warm-climate counterpart, especially France vs. Egypt (Pool et al., 2017).

### Fly rearing

Flies were reared at 25°C on Bloomington medium with a 12h:12h light:dark cycle. Larval density can impact cold tolerance (Henry, Renault, & Colinet, 2018), so when flies were to be used in experiments, egg density was standardised to approximately 50 per vial. Sexually mature females (presumed to be non-virgins) were isolated under CO_2_ on the day after emergence. CO_2_ exposure was brief (less than 10 minutes) and flies were then given a minimum of 48 hours to recover (Nilson, Sinclair, & Roberts, 2006). All flies used in experiments were between 5 and 7 days post-ecdysis.

### CCO

A fly’s chill coma onset temperature (CCO) is the temperature at which it becomes completely immobilized (Hazell & Bale, 2011; Overgaard & MacMillan, 2017). To measure this, we used a temperature ramping assay (Bertram, 1935; Sinclair, Coello Alvarado, & Ferguson, 2015). Briefly, the flies (eight to ten females per line) were placed individually in 3.7 mL glass screw-top bottles, which were clipped to a frame in a custom aquarium filled with a mix of ethylene glycol and water (1:1). The aquarium was connected to a 28 L programmable cooling bath (Model AP28R-30, VWR International, Radnor, USA) filled with the same ethylene glycol/water mixture. For the first 15 min of the experiment, the flies were held at their rearing temperature (25°C), and then the temperature was ramped down at 0.1°C/min. The temperature inside the aquarium was monitored with three type K thermocouples connected to a TC-08 recorder (Pico Technology, St Neots, UK), and the average of the three thermocouples was used to estimate fly body temperature. To test whether flies were still capable of movement at temperatures approaching the chill coma onset temperature, the vials and metal frame were periodically tapped with a metal rod, and the CCO was recorded as the temperature when this failed to stimulate any response.

### CCRT

Chill coma recovery time (CCRT) was measured by holding flies individually in 3.7 mL glass screw-top vials at 0°C (submerged in an ice-water slurry) for six hours, returning them to room temperature, and then measuring the time that it took for each one to stand on all six legs (MacMillan, Andersen, Davies, & Overgaard, 2015; Sinclair et al., 2015). Room temperature was recorded and accounted for in subsequent analysis. We measured eight to ten flies in most (26) lines, but in six, we measured between eleven and fifteen. In order to ensure a similar number of replicates in all lines, we randomly sampled ten replicates from each of those six lines (using the sample function in R) for the statistical analysis.

### Survival

Because of the large number of flies needed to accurately measure survival and the number of lines in our study system, and because this trait has already been characterised across a large number of lines in this system (Pool et al., 2017), only two populations were used in the survival experiment: France (5 lines total) and Egypt (4 lines). Flies were held in groups of 10 in vials (30 mL; containing c. 7 mL of medium) that were kept horizontal to reduce the risk of flies getting stuck in the food during exposure or recovery. The groups of flies were held at 4°C for zero to four days (4 to 9 replicates per line per day, where each replicate is a vial with 10 flies), given 24 hours to recover at 25°C, and then evaluated for survival using a 5 point scale (0: unmoving; 1: twitching but not standing; 3: standing but not walking; 4: walking but not climbing/flying; 5: climbing and/or flying) modified from MacMillan et al. (2018).

### Statistics

All analyses were performed in R version 3.6.3 (R Core Team, 2020). To examine the effect of population (e.g. the population derived from flies collected in the South African highlands) on the chill coma onset temperature (CCO), we fit a general linear model with CCO as the response variable and line nested within population as predictor variables. In all analyses, normality of the residuals was confirmed through visual inspection. We then used Tukey’s HSD (Honestly Significant Difference) test for pairwise comparisons among populations.

Likewise, to examine the effect of population on chill coma recovery time (CCRT), we fit a general linear model with the log transformed CCRT as the response variable (log transformation was necessary to ensure homogeneity of variance) with room temperature and line nested within population as predictor variables. Flies that did not recover during the 90 min observation time were given a CCRT of 90 minutes (17 out of 311 flies). Excluding these animals from the dataset entirely would artificially lower the mean, and we confirmed that doing so has no effect on our conclusions. Tukey’s HSD test was then used for pairwise comparisons among populations. To test for a correlation between CCO and CCRT, we calculated the mean CCO and CCRT of each line and fit a general linear model with mean CCO as the response variable, and population and mean CCRT as predictor variables. Because room temperature may affect CCRT, we also ran this analysis after adjusting CCRTs for the effect of room temperature extracted from the general linear model above. This adjustment, however, had no effect on our conclusion, so we opted to show the uncorrected data.

Mann-Whitney tests were used to compare population survival scores (which were not normally distributed) on each day of the chronic cold survival assay.

## Results

### Chill coma onset temperature (CCO) and chill coma recovery time (CCRT)

Population was a significant predictor of CCO (p < 0.001), but this was driven entirely by the Zambian population (our outgroup), which had a significantly lower CCO than all other populations (p < 0.001). There were no significant differences among the other populations (p > 0.05 in each pairwise comparison).

Similarly, population was a significant predictor of CCRT (p = 0.003), once fluctuations in room temperature were taken into account (ambient temperature has a significant impact on chill coma recovery time (p < 0.001)). However, in the pairwise comparisons, CCRTs differed significantly only between Zambia and both the French (p = 0.01) and Egyptian (p = 0.01) populations; there were no significant differences within population pairs (Figure 3). There was also no significant pattern in the number of flies that did not stand during the observation period (p > 0.05; Figure 3). Likewise, CCRT did not correlate with CCO (R^2^ = 0.00, p = 0.40), even when CCRTs were adjusted based on room temperature (R^2^ = 0.02, p = 0.30) (Figure 4).

**Figure 3.**
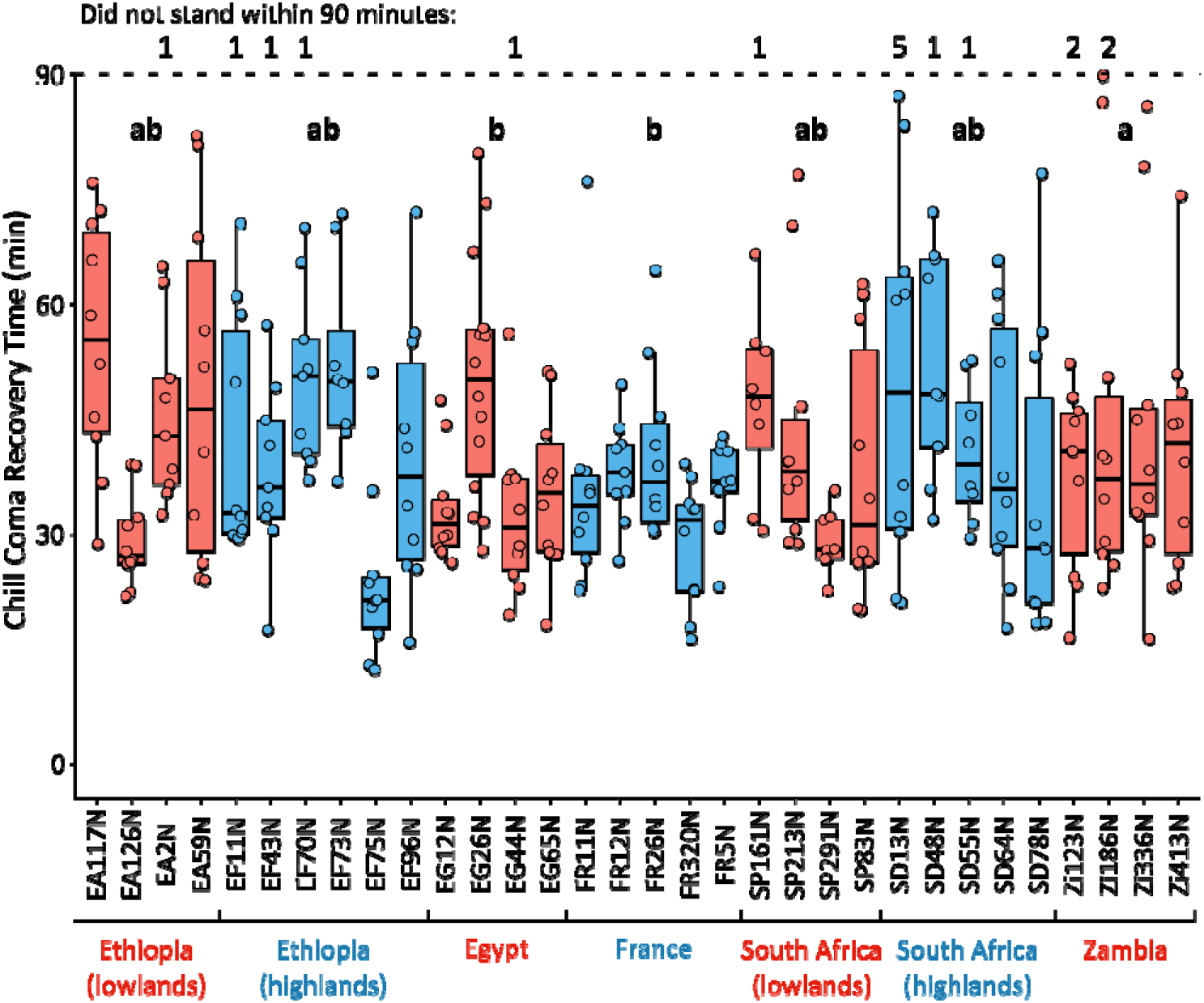
Chill coma recovery time of all lines. Lines in blue are from relatively cold habitats, whereas lines in red are from relatively warm habitats. The thicker line in the centre of each boxplot represents the median, and the upper and lower hinges represent first and third quartiles, respectively. Whiskers extend to 1.5 times the interquartile range, or to the smallest or largest values if they fall within 1.5 * IQR. Points represent individual flies. Different letters (a or b) indicate a significant difference (p < 0.05) at the population level.

**Figure 4.**
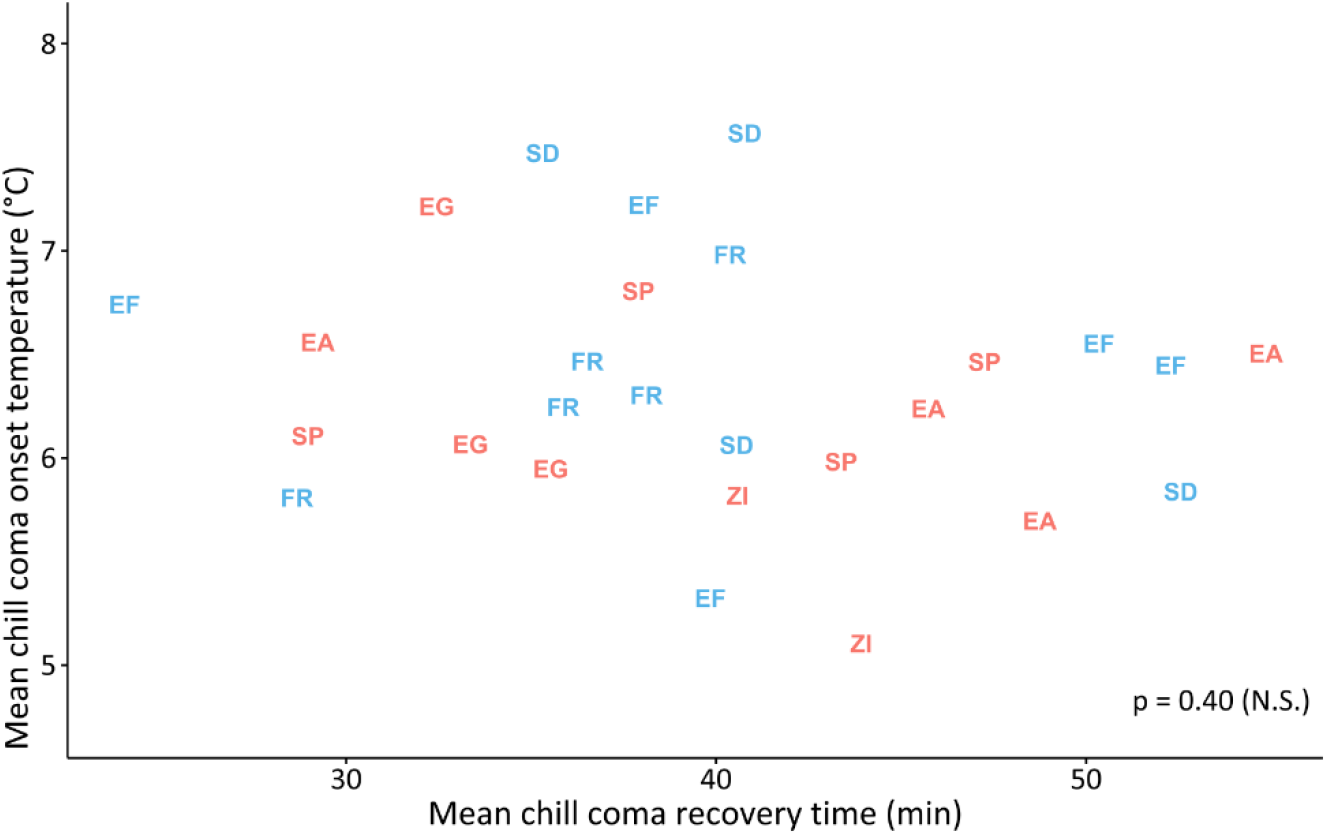
Mean chill coma onset temperatures (CCO) plotted against mean chill coma recovery times (CCRT) of all lines in all populations. Two-letter population codes are used as points to identify the population to which each line belongs: EA = Ethiopian lowlands; EF = Ethiopian highlands; EG = Egypt; FR = France; SP = South African lowlands; SD = South African highlands; ZI = Zambia. Colours indicate whether the climate of origin is relatively warm (red) or relatively cold (blue). We found no relationship between mean CCO and mean CCRT in the lines or populations studied.

### Survival

French lines had, on the whole, higher survival scores over time at 4°C than the Egyptian lines. The separation between populations was statistically significant on days one through four (p < 0.001) and was largest after two days at 4°C (Figure 5).

**Figure 5.**
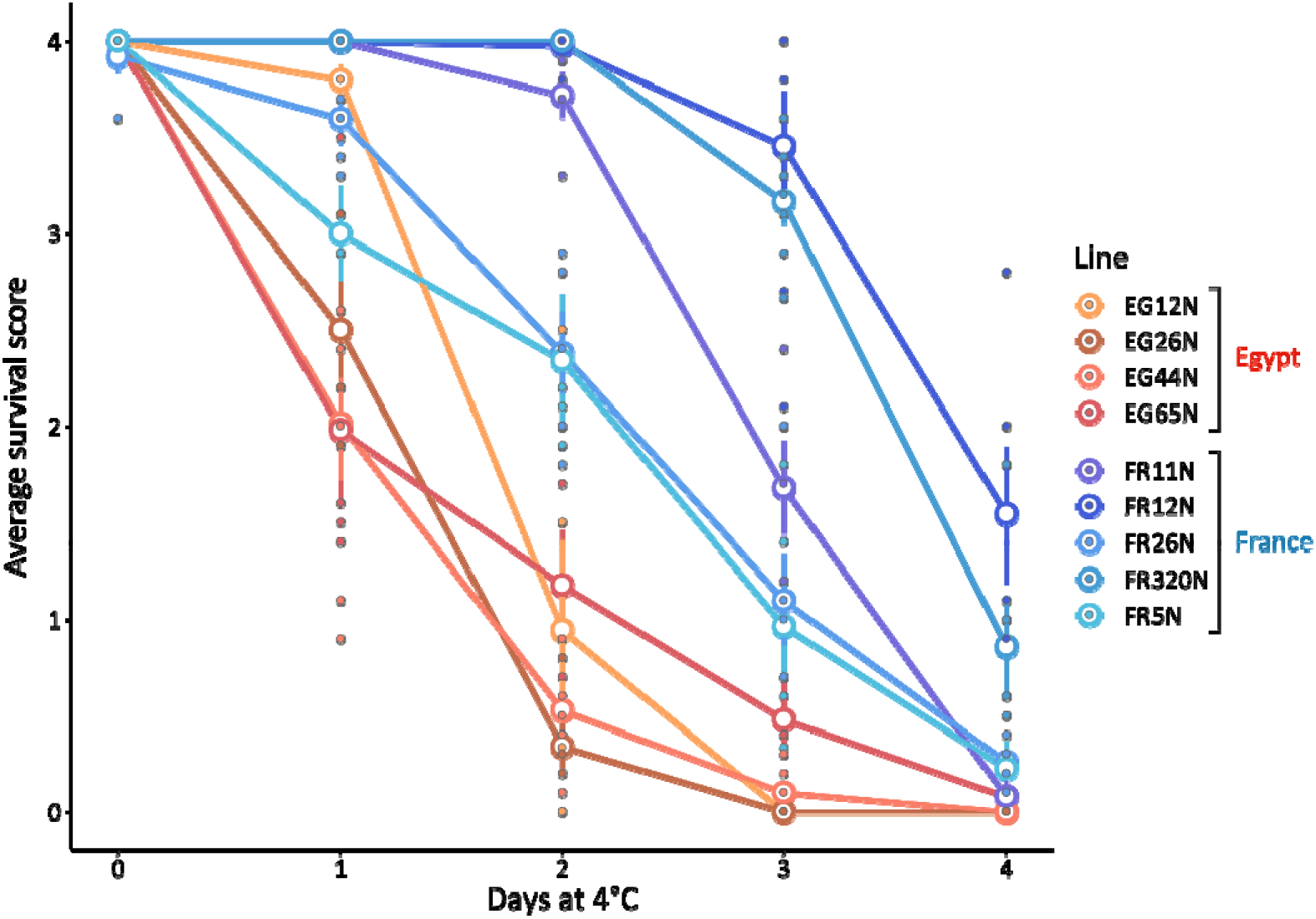
Average survival score after exposure to 4°C for between zero and four days for four Egyptian lines and five French lines. Each point represents a single replicate: the average survival score of a vial of approximately ten flies. A survival score of zero meant that a fly was not moving, whereas a survival score of four meant it was climbing and/or flying. Survival scores were recorded 24 hours after removal from the cold (or 24 hours after the beginning of the experiment, in the case of flies that were not exposed to 0°C). Open circles represent line means, and error bars represent standard errors. Some error bars are too small to be visible.

## Discussion

In this study, we confirmed that chill tolerance traits can vary independently in *Drosophila melanogaster*. Similar results have emerged in laboratory selection experiments (Garcia et al., 2020; Gerken et al., 2016) where selection pressure is controlled, but never, to our knowledge, in wild-derived populations like those used here. We were surprised to find that this was the case even in populations that are known to differ significantly in one chill tolerance trait (Pool et al., 2017) – in a way that matched the climatic conditions where their ancestors were collected. In some ways, this finding is not surprising: it is increasingly clear that the underlying mechanisms of several commonly measured chill tolerance traits differ (M. K. Andersen & Overgaard, 2019; Overgaard & MacMillan, 2017), and it is also possible to select on such traits separately (Garcia et al., 2020; Gerken et al., 2016). However, it remains unclear why some wild-derived populations from colder regions display greater chill tolerance across a range of traits (A. A. Hoffmann et al., 2005; Ary A. Hoffmann et al., 2002; Overgaard et al., 2011), whereas others (e.g. the populations studied here) can only be differentiated if we measure the “right” trait. In other words, in order to draw meaningful conclusions about differences in stress tolerance (or the lack thereof), we strongly advise researchers measure more than one trait.

There are several possibilities for why some cold climate populations might not evolve improvements in specific chill tolerance traits. First, it is possible that after evolving better chill tolerance in one trait, or other adaptations to a cold climate that we were unable to detect, these populations did not face the necessary selection pressure to drive improvements in other traits. Notably, however, at an interspecific level this does not appear to be the case: *Drosophila* species from cold climates do not evolve one chill tolerance trait and then stop (J. L. Andersen et al., 2015). Second, there may not have been enough evolutionary time for the populations to fully differentiate: *D. melanogaster* only left Sub-Saharan Africa some ten thousand years ago (Baudry, Viginier, & Veuille, 2004), and it is possible that some chill tolerance traits may respond more quickly to the selection pressures of a cold climate than others. However, Australian populations have been shown to differ across multiple chill tolerance traits (A. A. Hoffmann et al., 2005; Ary A. Hoffmann et al., 2002; Overgaard et al., 2011) – and the species only reached that continent within the past few hundred years (Baudry et al., 2004). Finally, populations that do not appear to differ based on multiple chill tolerance traits may differ in their plastic responses to cold. Adult and developmental cold acclimation improve multiple chill tolerance traits (e.g. reducing chill coma onset temperatures, increasing survival after cold shock, etc.) (MacMillan, Andersen, Loeschcke, et al., 2015). Plastic responses can be very strong and may be more ecologically important than differences in basal (i.e. non-plastic) chill tolerance (A. A. Hoffmann et al., 2005).

We note that the one trait that distinguished our populations, survival after 4 days at 4°C, was one that may have permitted plasticity (specifically acclimation) during the assay. When held at temperatures slightly below the chill coma onset temperature, some *Drosophila* species, including *D. melanogaster*, show evidence of acclimation by recovering the ability to stand shortly after falling into a chill coma (MacMillan, Andersen, Davies, et al., 2015) or displaying improved cold shock resistance after exposure to 4°C for several days (Chen & Walker, 1994). Differences in chill tolerance plasticity have been studied on an intraspecific level before: Australian *D. melanogaster* populations, which have been shown to differ in multiple chill tolerance traits along a latitudinal gradient, do not differ in their response to either adult or developmental acclimation treatments (Ary A. Hoffmann & Watson, 1993). However, a much larger study of *D. melanogaster* populations from around the world, including Africa and Europe, found that flies from temperate regions do indeed respond more strongly to a developmental acclimation treatment (i.e. their chill tolerance improved more) than flies from tropical regions (Ayrinhac et al., 2004). That study measured only one trait, CCRT, and also found that flies from colder regions tended to have lower CCRTs than flies from warmer regions – a pattern that we did not observe in our study populations – but it demonstrates that intraspecific variation in the capacity for acclimation is possible.

It remains to be shown whether plasticity is behind the discrepancies that we observed – we saw essentially no differentiation in two commonly- and rapidly-measured traits (CCO and CCRT), and clear differentiation in one trait (survival after 4 days at 4°C) measured under conditions that would normally induce some form of acclimation. Nevertheless, we feel that this system, which contains populations that adapted separately to cold environments in Africa and Europe, would lend itself well to future studies of how different forms of plasticity affect different thermal tolerance traits. It may also offer an opportunity to explore potential trade-offs between basal and plastic cold tolerance, which have previously been explored in drosophilids (Gerken, Eller, Hahn, & Morgan, 2015; Nyamukondiwa, Terblanche, Marshall, & Sinclair, 2011) (but see also van Heerwaarden and Kellermann, 2020, who call previous work into question and discuss the challenges of accurately identifying basal vs. plastic thermal tolerance trade-offs).

An important caveat here is that we worked with laboratory-reared flies. However, past research has shown that chill tolerance traits are remarkably stable in lab reared flies over time (Ayrinhac et al., 2004; Gilchrist, Huey, & Partridge, 1997; Maclean, Kristensen, Sørensen, & Overgaard, 2018), and since we were able to confirm that the large difference in chill tolerance previously observed by Pool et al. (2017) between the French and Egyptian populations still exists. Therefore, we are confident that differences in chill tolerance seen in these populations reflect differences in their wild progenitors and are worthy of further study.

## Conclusion

A cold environment can exert selection pressure on an insect and impact diverse chill tolerance traits in unexpected ways. To our surprise, traits that would allow higher activity in the cold (i.e. a low chill coma onset temperature) or faster recovery from chill coma have apparently not been selected for in the cold-climate African and European *D. melanogaster* populations tested here – but the ability to survive a chronic, mild cold stress has. These findings support the notion that chill tolerance traits are mediated by different physiological mechanisms and highlight the importance of measuring more than one chill tolerance trait in wild-derived populations. A benefit of such paired population study systems is that they can allow us to see how chill tolerance and its underlying mechanisms can evolve under similar low temperature selection pressures.

## Supporting information

Data archive for Davis et al.

## Acknowledgements

The authors wish to thank Erica O’Neill and Yingtong Gao for their assistance tipping flies during the time this research was being conducted, as well as the whole MacMillan Lab for providing feedback on an early draft of this manuscript. This work was supported by a Natural Sciences and Engineering Research Council (NSERC) Discovery Grant to H.A.M. (RGPIN-2018-05322), an NSERC Canada Graduate Scholarship to H.E.D., and infrastructure funding to H.A.M. from the Canadian Foundation for Innovation and Ontario Research Fund Small Infrastructure Fund.

